# Partner fidelity with root symbionts impacts plant performance in the face of changing above- and belowground community context

**DOI:** 10.64898/2026.01.13.699266

**Authors:** Anne M. McLeod, Daniel B. Stouffer, Jason M. Tylianakis, Warwick J. Allen, Lauren P. Waller, Ian A. Dickie, Hao Ran Lai, Bernat Bramon Mora, Jonathan D. Tonkin

## Abstract

Global environmental changes generate novel communities. Some species adapt to novel community contexts by demonstrating interaction fidelity, or consistently interacting with the same species, while others demonstrate interaction flexibility, or interacting with whichever partners are available (i.e., rewiring). However, the drivers and benefits of fidelity are unclear. Here, we use data from a large-scale mesocosm experiment to determine whether plant characteristics (e.g., provenance, functional group) and community contexts impact plant interaction fidelity to belowground mutualist and antagonist partners (i.e., root fungi and bacteria), and how this fidelity relates to plant performance. We found that the plant-antagonist relationships exhibited higher fidelity than plant-mutualist relationships with the strength of fidelity depending on a combination of plant characteristics and community context. Conversely, plant-mutualist fidelity impacted plant performance with competing effects depending on plant provenance and aboveground herbivore presence. Our study advances understanding of how species interactions influence plant performance in a changing world.

**Statement of authorship:** AMM, JDT, JMT and DBS designed the concept. All authors contributed to refining the ideas. AM coded the analysis to produce the results with contributions from JDT, DBS, and BBM. HRL and DBS helped AMM with the statistical analysis. LPW, WJA, IAD, and JMT designed the experiment while LPW and WJA conducted the experiment. AMM wrote the first draft of the manuscript with assistance from all authors. IAD, JMT, and JDT obtained funding. All authors contributed to further editing of the manuscript.

## Introduction

As anthropogenic change disrupts communities, species can either remain faithful to, or be flexible in, their interspecific interaction partners (CaraDonna *et al*. 2017; Spiesman & Gratton 2016). Although multiple studies predict that higher interaction flexibility (also called ‘rewiring’) increases the robustness of species interaction networks to species extinctions (e.g., Kaiser Bunbury *et al*. 2010; Ramos Jiliberto *et al*. 2012; Valdovinos *et al*. 2013), such flexibility may come with population-level trade-offs for the species involved, exemplified by empirical evidence for the conservation of species roles across a wide range of antagonistic and mutualistic networks (Emer *et al*. 2016; Rezende *et al*. 2007; Stouffer *et al*. 2012). There likely exists a gradient whereby, as species’ ranges begin to shift and novel communities form, it may be advantageous to remain faithful to interaction partners. For example, species may specialise on key mutualist partners, with whom they interact more consistently through time and achieve greater fitness benefits (Peralta *et al*. 2020). As the community changes further, however, interaction flexibility is likely increasingly important. This may be especially true when an individual is pushed beyond its ‘biotic comfort zone’, as flexibility better allows individuals and populations to partition niches and minimise realised competition (Pianka 1974; Spiesman & Gratton 2016). Thus, the exact circumstances under which remaining faithful to interaction partners is a strategic benefit or comes at a cost remain unclear.

Conservation of interspecific interaction partners can be measured as partner fidelity (i.e., the tendency for two species to interact everywhere they co-occur) (Fortuna *et al*. 2020). Partner fidelity can be measured across both antagonistic interactions and mutualistic interactions (Morris *et al*. 2007); however, the costs and benefits of partner fidelity likely depend on the type of interaction. For example, it may be more advantageous for species to interact consistently with the same beneficial, mutualistic partners than the same detrimental, antagonistic partners, either because specialised enemies such as pathogens may cause more damage to their hosts due to the evolution of effective countermeasures to the chemical defenses of plants (Berenbaum & Zangerl 1998) or because traits involved in antagonistic interactions are more evolutionarily malleable than those linked to mutualistic interactions (Fortuna *et al*. 2020). Understanding these trade-offs between mutualisms and antagonisms is challenging, since research typically focuses on a single type of interaction partner (Porter *et al*. 2020). This challenge is further complicated by evidence that the strengths of different types of biotic interactions are interdependent (Morris *et al*. 2007; Sauve *et al*. 2016), meaning that apparent relationships involving one interaction type could arise through its covariation with another. For example, both herbivores and plant pathogens can select for generalised defense compounds in plants which in turn inhibit both types of enemies (Biere *et al*. 2004).

Plants engage in a complex network of interactions with their mutualist and antagonist partners. These interactions are often highly plant species-specific, including preferentially interacting with different species of mycorrhizal fungi (Martínez García *et al*. 2015; Šmilauer *et al*. 2020; Vályi *et al*. 2015), potentially related to functional differences of the plants. For example, forbs benefit more from their arbuscular mycorrhizal (AMF) partners than grasses (Hoeksema *et al*. 2010; Wilson & Hartnett 1998), while more specialised ectomycorrhizal fungi interact primarily with woody plants (Smith & Read 2010). Moreover, it has been demonstrated that AMF diversity is lower in communities dominated by non-native plants as non-native plants promote general fungal partners (Ramana *et al*. 2025). Thus, plant interaction fidelity may be expected to change depending on several factors, including plant species characteristics (e.g., provenance, functional group).

Plant interaction fidelity may also be expected to change depending on local community context. Competition with other plants has been shown to decrease the efficacy of host plant defenses (Seabloom *et al*. 2018), perhaps resulting in lower antagonistic partner fidelity in highly competitive environments. In contrast, plant interaction fidelity for AMF can be consistent across soils with different edaphic conditions and inoculum pools (Lewe *et al*. 2025), suggesting that fidelity to fungal mutualists may depend less on environmental context. To date, however, most studies that examine interaction flexibility have used observational data collected along spatial or temporal gradients (McLeod *et al*. 2020; Trøjelsgaard *et al*. 2015). This makes it challenging to disentangle the impact of community context from environmental change, particularly because observational studies may contain a reduced species pool due to environmental filtering. Instead, we need experimental studies that manipulate broad biotic gradients to explicitly test how species and their interactions with mutualists and antagonists respond to being pushed beyond their biotic comfort zone – a critical step to improving our ability to predict how interspecific interactions mediate plant performance in a changing world.

Here, we study data from a large-scale experimental multitrophic mesocosm system to investigate how plant characteristics and changing community contexts impact individual plant interaction fidelity to belowground mutualist and antagonist partners, and how this in turn influences plant performance (i.e., biomass). We directly manipulated both competitive plant-plant interactions and herbivory by changing plant community composition and the presence of invertebrate herbivores to assess the indirect impact of aboveground community on belowground partner fidelity. We also manipulated the soil microbial communities to directly impact the assemblage of belowground interacting partners – characterised using DNA-metabarcoding of root tissue samples and used to measure partner fidelity of each individual plant (henceforth individual plant interaction fidelity) for each type of interaction partner (bacterial antagonist, fungal antagonist, fungal mutualist). We then used these data to test four main hypotheses (see Appendix S1 for further description):

(H1) We hypothesised that individual plant interaction fidelity would depend on the type of interaction partner and be highest for fungal mutualists, and that individual plant interaction fidelity to different types of interaction partners would be interdependent.

(H2) We hypothesised that plant characteristics would influence individual plant interaction fidelity and that the direction of the effect would depend on the type of interaction and plant characteristic (e.g., non-native plant species should exhibit lower plant interaction fidelity to fungal mutualists than native plant species).

(H3) We hypothesised that community context (the functional groups and provenance of other plant species in the community) would influence individual plant interaction fidelity and that the direction of the effect would depend on the type of interaction and plant characteristic, driven mostly by competition.

(H4) We hypothesised that the cost of high interaction fidelity will depend on the interaction type. High interaction fidelity to antagonists is hypothesized to be costly for individual plants as more specialised enemies are often more damaging, while high interaction fidelity to mutualists, mean they will be unable to adapt to the challenges of a changing community and thus high interaction fidelity will lead to reduced plant performance.

## Methods

We used 160 mesocosms to manipulate grassland plant communities (for more details see Allen *et al*. 2021; Waller *et al*. 2020). From a pool of 39 total plant species, we assembled 20 unique communities, comprising one plant individual from each of 8 plant species, with 8 replicates of each community. Each plant species was categorised as either native (plant species that co-occur in New Zealand grassland communities, 19 species) or non-native (plants which have been introduced to New Zealand grassland communities, 20 species). Plant species were also categorised according to whether they were grasses (11 species), woody species (15 species), or forb species (13 species) and then whether they were legumes (8 species). Twenty unique plant communities were designed based on their percentages of non-native and woody species (varying orthogonally from 0 – 100% and 0 – 63%, respectively, with non-woody species consisting of both forbs and grasses). This experimental design resulted in gradients of non-native, legume, woody, forb and grass species, allowing us to assess the effect of plant community context on individual plant interaction fidelity with their belowground interaction partners. We further manipulated the community by imposing an herbivory treatment (added, removed), and soil treatment (home, away), which were applied factorially across the eight replicates per community (see Appendix S2 and Waller *et al*. 2020, 2024).

Individual plants were grown from seeds or cuttings for one month before being transplanted into mesocosms. After a year, individual plants were harvested from the experimental mesocosms, above and belowground dry biomass were measured, and roots were processed for molecular sequencing of the bacterial and fungal associations. In total, there were 6 – 52 individuals of each species across the 160 mesocosms both because of the design of the communities and plant mortality.

### Molecular Sequencing

DNA was extracted from a total of 895 roots (i.e., the number of living plants at the end of the experiment) harvested from the 160 mesocosms at the end of the experiment using MoBio PowerSoil (QIAGEN) extraction kits (full description can be found in Waller et al. 2024). We characterised the fungal and bacterial communities by amplifying the internal transcribed spacer (ITS) of the ribosomal RNA (rRNA) operon for fungi and 16S rRNA for bacteria using polymerase chain reaction (PCR) with barcoded primers fITS7/ITS4 (Ihrmark *et al*. 2012) and 515FB/806RB, respectively. Amplicons were sequenced on an Illumina MiSeq analyser using the 600-cycle Reagent Kit V3, delivering 2 X 300 base pair reads/sequence (Illumina, San Diego, California, USA).

Sequences were paired and chimeras were removed, then clustered into operational taxonomic units (OTUs) at 97% sequence similarity using Vsearch (Rognes *et al*. 2016). The taxonomic database FUNGuild (Nguyen *et al*. 2016) was used to assign functional attributes to the fungal taxa, and we retained only the taxa assigned as “probable” or “highly probable” mutualists and pathotrophs for subsequent analysis. Mutualists included OTUs returned as “arbuscular mycorrhizal” and “ectomycorrhizal”. We excluded antagonistic taxa with flexible strategies, (i.e., “pathotroph-saprotroph”), as we were only interested in antagonists that harm live host cells (i.e., biotrophic, rather than necrotrophic pathogens). The functional attributes of bacteria were assigned based on information from Makiola *et al*. (2019) and information provided in Martinez-Romero *et al*. (2024). We did not consider bacterial mutualists (e.g., rhizobia, *Frankia*) due to the complexity of mapping symbioses within the Rhizobiales (Wang *et al*. 2020) and the lack of sufficient information on New Zealand strains using modern molecular techniques.

There has been much debate about how to treat low abundance OTUs, because clustering programs can generate many spurious, low abundance OTUs, which are the result of PCR bias and technical errors resulting in an artificially high number of singletons (i.e., OTUs observed only once; Lindahl *et al*. 2013; Medinger *et al*. 2010; Reeder & Knight 2009; Tedersoo *et al*. 2010). This was unlikely to influence our results, because singletons were necessarily removed during the calculation of partner fidelity (see next section). Thus, the OTUs included in our hypothesis tests were likely present in the plant roots and not an artefact.

### Measuring Interaction Fidelity

We compared a focal plant individual’s locally observed interactions with the interactions observed for conspecific plant individuals found in other mesocosms. We quantified this using the focal individual’s average Sorensen similarity across pairwise comparisons (e.g., Koleff *et al*. 2003; Poisot *et al*. 2012). That is, for every individual plant (*P_iA_* where *i* is the plant species and *A* is the mesocosm in which that individual plant is found) we averaged the *N_i_*-1 Sorenson similarities across mesocosms (where *N_i_* is the number of mesocosms in which focal plants of species *i* are found; see Fig. 1). Note that we considered only interactions with the pool of shared bacterial or fungal species in each mesocosm pair. This ensured that the absence of an interaction between a plant and a bacterial or fungal partner was a true absence of the interaction (given co-occurrence of the species) and not an artefact of the bacterial or fungal species not being locally available (Poisot *et al*. 2012). Sorenson similarities range from 0 to 1, where 0 indicates no overlap in interaction partners and 1 indicates complete overlap (i.e., that individuals of a plant have complete fidelity to the same interaction partners wherever they co-occur).

**Fig. 1.**
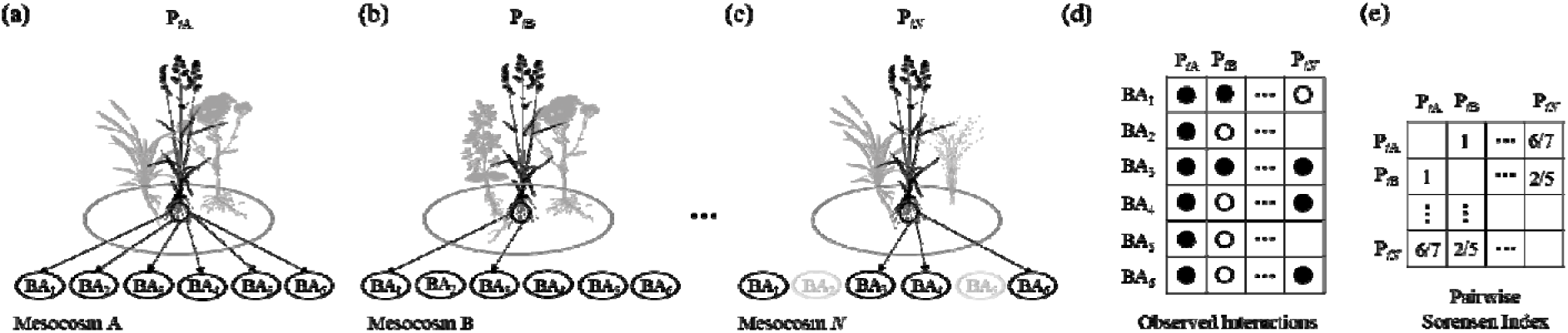
Overview of calculating individual plant interaction fidelity to bacterial antagonist partners. (a - c) Here, we depict hypothetical mesocosms *A*, *B*, through to *N_i_* where *N* is the total number of mesocosms in which focal plants of species *i* are found. Across the *N_i_*mesocosms, focal plants of species *i* interact with six different bacterial antagonist species (denoted BA*_j_* where *j* is the hypothetical species number). (a) All six bacterial antagonist species are found in the mesocosm A (denoted with the black ovals), and focal plant P*_iA_* interacts with all six (denoted with arrows). (b) In mesocosm B, focal plant P*_iB_* only interacts with two of the bacterial antagonists (BA_1_ and BA_3_), despite all six bacterial antagonist species being locally present. (c) BA_2_ and BA_5_ are not locally observed in mesocosm *N* (denoted with the grey circle), and focal plant P*_iN_*only interacts with three of the four locally observed bacterial antagonist species. (d) These mesocosm level plant-bacteria interactions can be turned into a matrix of interactions between focal plant individuals and each bacterial antagonist species, BA*_j_*, where empty circles represent the presence of the bacterial antagonist species in the mesocosm (but no interaction) and filled circles represent an observed interaction between the focal plant individual and bacterial antagonist species BA*_j_*. To measure the interaction fidelity of each individual plant, we compare the interactions for that individual to the interactions of individuals from the same plant species found in all other mesocosms, considering only those bacterial antagonist species that are shared between the two mesocosms. (e) Using these values, we calculate the pairwise Sorensen similarity index, e.g., pairwise Sorensen index between P*_i_*_A_ and P*_iN_* is 6/7 given there are 3 shared interactions and one additional interaction in mesocosm A with a species also present in mesocosm N (pairwise Sorensen index is two times the number of shared interactions divided by two times the number of shared interactions plus the number of interactions unique to a single mesocosm for a species found in both mesocosms), resulting in *N_i_*-1 Sorensen similarities for each mesocosm depicted in the matrix shown in (e).

We wanted to ensure that values of our averaged Sorensen similarity index were meaningful indicators of interaction fidelity and not, for example, high on average due to a plant individual always interacting with the full suite of interaction partners available to it (i.e., if only one potential partner species is ever available, all individuals from a given plant species may always seem to have high interaction fidelity). We thus compared our observed Sorensen index values to those obtained using a null model that randomised the interactions within each network (see Fig. 1). To do this, we used the total pool of fungi and bacteria in a mesocosm to randomly select partners for each plant individual in that mesocosm, while fixing the plant individual’s degree (i.e., number of interaction partners) for each interaction partner type. In that way, if individual plant *P_iA_* has four fungal mutualist partners, four possible partners were selected at random and without replacement from the pool of all fungal mutualists in mesocosm *A*. We did this for all individuals of a given plant species (each in a different mesocosm) and then calculated Sorensen similarity indices as described above. We repeated this null model randomization 1000 times, then calculated the null average Sorensen similarity index for each plant individual across the 1000 draws. For each plant individual in each mesocosm, we then calculated individual plant interaction fidelity as the z-score from the comparison of their observed Sorensen similarity index against the null model distribution (see Fortuna *et al*. 2020). This process produced a measure of individual plant interaction fidelity that was scaled according to the potential range of interaction fidelity values that could occur at random (see Appendix S2 for more information on how to interpret the value). We repeated the null model comparison separately for each type of interaction partner: fungal mutualist, fungal antagonist, and bacterial antagonist.

We removed three of the original 39 plant species which were only present in, at most, two mesocosms by the end of the experiment and thus lacked sufficient variation to compare against the null model results (Appendix S2, Table S1).

### Statistical Analysis

First, we used linear mixed models (Gaussian family and identity link function) to determine the impact of species characteristics and community context on each type of plant interaction fidelity (i.e., plant interaction fidelity to bacterial antagonists, plant interaction fidelity to fungal antagonists, and plant interaction fidelity to fungal mutualists). The general model structure for each type of interaction fidelity was the same and included main effects of species’ normalised degree for a given type of interaction partner (i.e., a measure of generalism; further defined in Appendix S2), whether the species was woody, grass, forb, legume, or non-native, the community context (i.e., percentage woody, grasses, forbs, legumes, and non-native plants in the mesocosm community), soil treatment (home, away), herbivore treatment (added, removed), and then five statistical interactions between plant characteristics and community context (i.e., whether the plant was woody/grass/forb/legume/non-native and the corresponding percentage of woody/grass/forb/legume/non-native in the mesocosm community). Mesocosm community (i.e., the 20 different plant communities) was included as a random intercept, and nested within that, mesocosm identity to control for non-independence of plants in the same mesocosm. We constructed our models based on our key hypotheses, thus trying to keep them simple enough to interpret, but also to include statistical interactions for which we had clear hypotheses.

To understand the impact of interaction fidelity on plant biomass, we again used a linear mixed model (Gaussian family and identity link function with log biomass as the response variable to meet assumptions of normality) that included main effects of individual plant normalised degree for each type of interaction partner, individual plant interaction fidelity for each type of interaction partner, whether the species was woody, grass, forb, legume, or non-native, the community context (i.e., percentage woody, etc. in the mesocosm community), soil treatment, herbivory treatment, and then the same five statistical interactions between plant characteristics and community context as above. We further included (1) the statistical interaction between plant provenance and the herbivory treatment as a previous study found it to be a meaningful predictor of plant biomass (Allen *et al*. 2021) and (2) the statistical interactions between each type of interaction fidelity (e.g., the statistical interaction between individual plant interaction fidelity to fungal mutualists and interaction fidelity to fungal antagonists). Additionally, we included the statistical interactions between each type of individual plant interaction fidelity and each of the other main effects (e.g., percentage of woody species in the mesocosm community) to test if the influence of individual plant interaction fidelity was context-dependent In addition to the random effects of mesocosm community and mesocosm identity (as above), we included plant species as a random intercept to account for differences in average biomass associated with each plant species. The analysis was done with both total plant biomass and belowground biomass as the response variable because plants’ belowground biomass has been shown to be impacted by availability of resources (Bloom *et al*. 1985; Chen & Reynolds 1997; Poorter *et al*. 2012). For more information on how each predictor was calculated see Appendix S2.

## Results

Counter to H1, individual plants showed higher interaction fidelity to bacterial antagonists and fungal antagonists than expected by chance, as individual interaction fidelity was greater than zero for all but two of the 815 plant individuals (Fig. 2a and b). In contrast, only 19 out of the 36 plant species had individual plant fidelities to their fungal mutualists which never overlap with zero, with 27 plant individuals showing individual interaction fidelity less than zero (Fig. 2c). No plant species showed lower average interaction fidelity than expected by chance for any type of interaction partner.

**Fig. 2.**
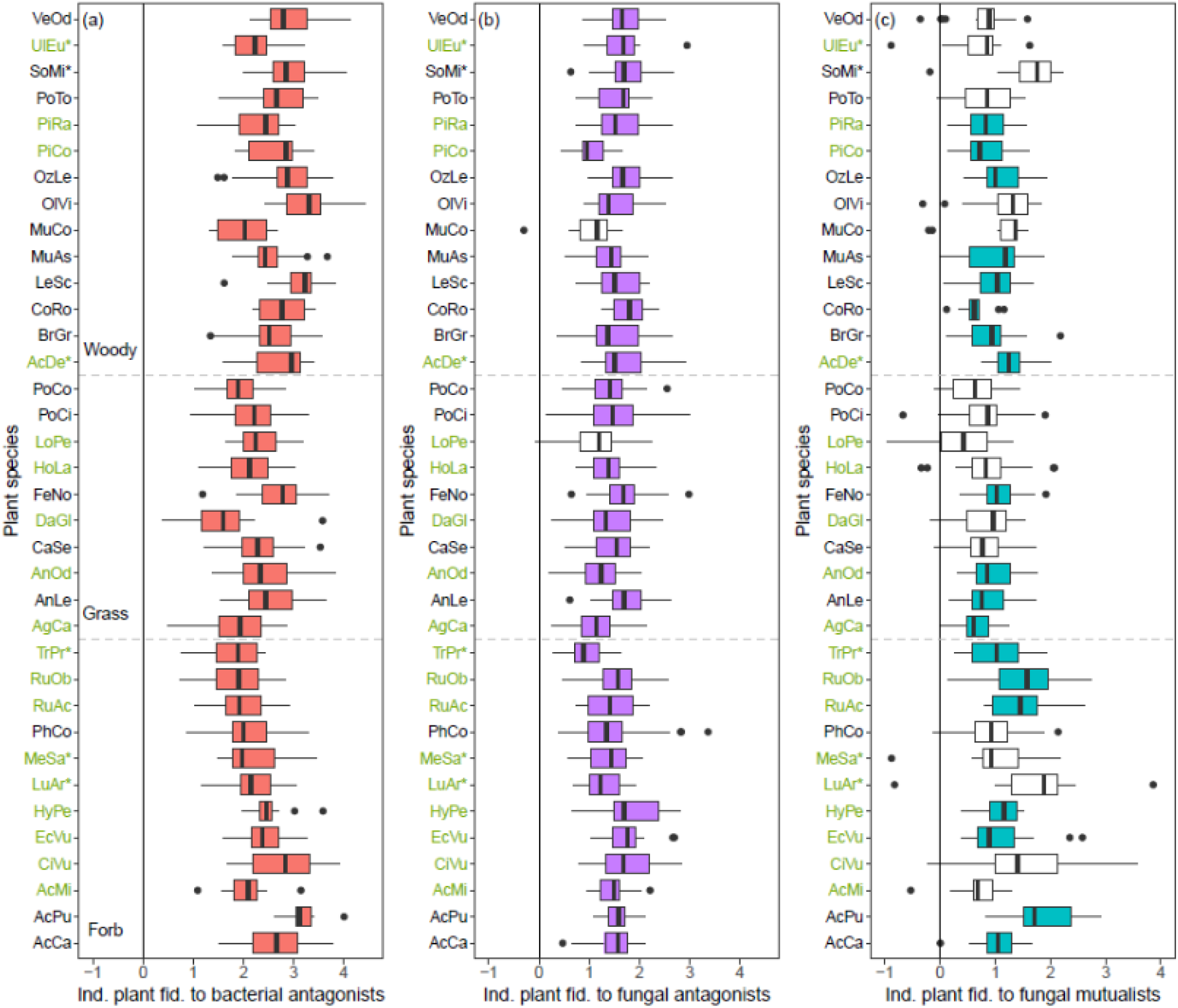
Plant interaction fidelity to (a) bacterial antagonist partners, (b) fungal antagonist partners, and (c) fungal mutualist partners for individuals from each of the 36 plant species. Plant interaction fidelity is measured as the z-scores (i.e., deviation from the null model of random interactions), summarised by boxes that span the 25^th^ – 75^th^ percentile and whiskers that extend to the largest value no further than 1.5 × the interquartile range. Boxes are coloured for plant species whose individual plant fidelities never overlap with zero (vertical solid line). Each species is identified by a four-letter code along the y-axis (see Appendix S2 Table S1 to match the 36 plant species to the codes). Woody (top), Grass (middle), and Forb (bottom) plant species are separated by horizontal dashed lines. Legumes are denoted with an * beside their four-letter code. Non-native species are denoted with green text.

Individual plant interaction fidelity to fungal mutualists and fungal antagonists had a strong positive correlation (H1; Table 1 and Fig. 3a, turquoise line). Individual plant interaction fidelity to bacterial antagonists was also positively correlated with interaction fidelity to fungal antagonists (Table 1 and Fig. 3a red line).

**Fig. 3.**
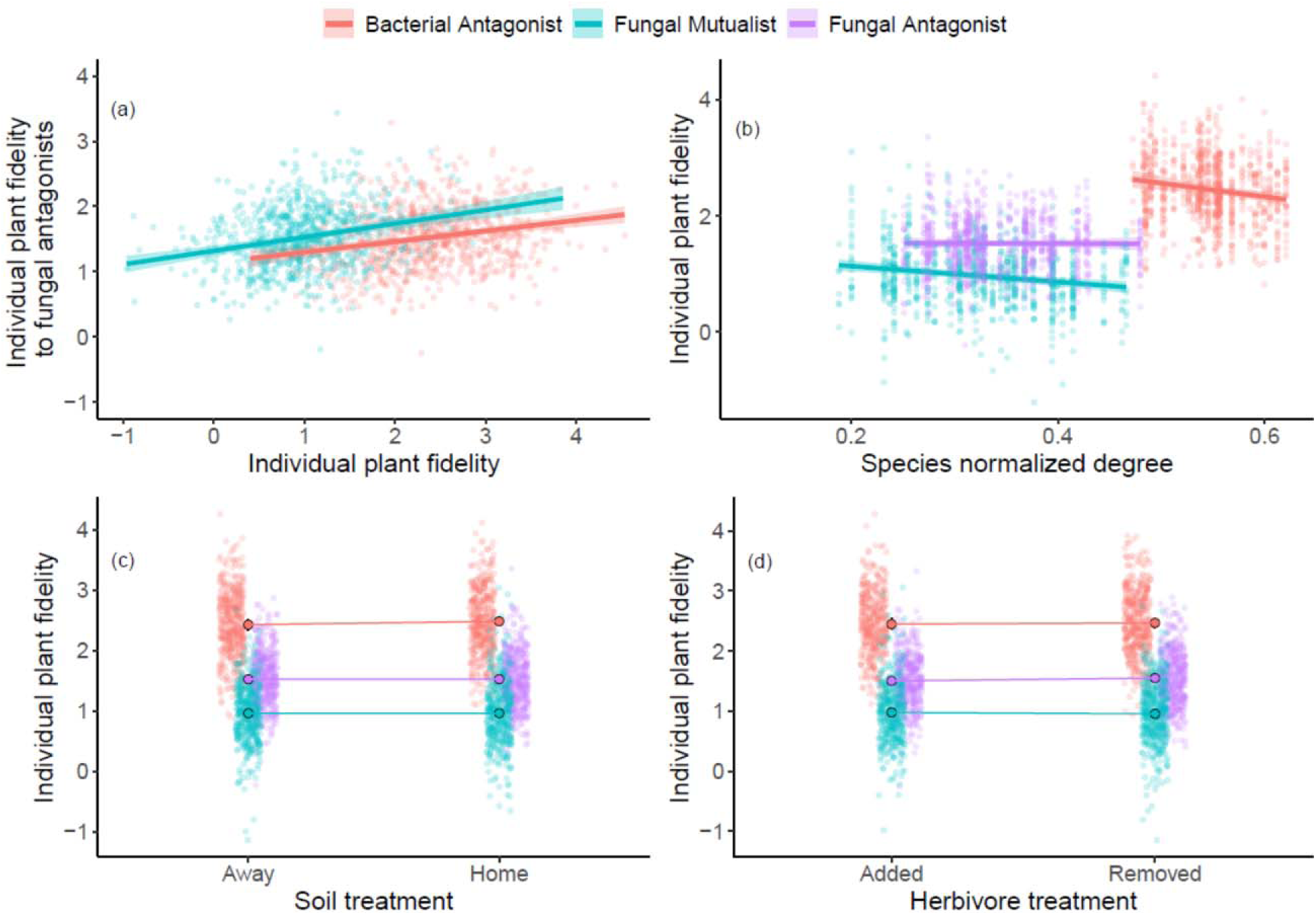
The inferred relationships between (a) individual plant interaction fidelity to fungal antagonists and their individual plant interaction fidelity to either bacterial antagonists (red) or fungal mutualists (turquoise), and (b-d) individual plant interaction fidelity to bacterial antagonist (red), fungal mutualist (turquoise) and fungal antagonist (purple) partners’ responses to (b) the corresponding species’ normalised degree, (c) soil treatment, and (d) herbivore treatment ± one standard deviation (shaded area and error bars – although error bars are very small in (c) and (d)). See Table 1 for details on statistical significance. Note that non-focal predictors were held at their mean values and all lines were fit using the linear mixed models described in the text.

**Table 1.** Relationship between species characteristics and mesocosm context with individual plant interaction fidelity to bacterial antagonists, fungal antagonists, and fungal mutualists. Here, Fid. is fidelity, sp. is species, Sp. normalised degree is the average number of interaction partners a plant from that species has with a given partner type (bacterial antagonist, fungal antagonist, or fungal mutualist) adjusted by the total richness of that partner type in the mesocosm, Woody (%) is the percentage of woody species in that mesocosm (not counting the focal plant), and the same interpretation applies for Grass (%), Legume (%), and Non-native (%). Legume sp. is whether or not the focal plant is a legume, Woody sp. is whether or not the focal plant is woody, Grass sp. is whether or not the focal plant is a grass, Non-native sp. is whether or not the focal plant is non-native, Soil trt. is the soil treatment (home or away), and Herb trt. is the herbivore treatment (added or removed). Bold values represent significant relationships. Marg. R^2^ and Cond. R^2^ represent marginal and conditional R^2^, respectively, where conditional R^2^ accounts for random effects. Note, explicit forb species are absent from the table because, by deduction, they correspond to species that are neither woody nor grass species.

| *Predictors* | *Fid. to Bacterial* |  | *Fid. to Fungal* |  | *Fid. to Fungal* |  |
| --- | --- | --- | --- | --- | --- | --- |
|  | *Antagonists* |  | *Antagonists* |  | *Mutualists* |  |
|  | *Est.* | *Std. Error* | *Est.* | *Std. Error* | *Est.* | *Std. Error* |
| Fid. to fungal mutualists |  |  | **0.236** | **0.034** |  |  |
| Fid. to bacterial antagonists |  |  | **0.207** | **0.035** |  |  |
| Sp. normalised degree | **-0.138** | **0.043** | -0.004 | 0.041 | **-0.157** | **0.034** |
| Legume sp. | -0.114 | 0.138 | -0.001 | 0.142 | **0.342** | **0.144** |
| Woody sp. | **0.510** | **0.079** | 0.056 | 0.078 | **-0.235** | **0.077** |
| Grass sp. | -0.026 | 0.093 | 0.064 | 0.09 | **-0.441** | **0.085** |
| Non-native sp. | -0.017 | 0.089 | 0.134 | 0.083 | **0.274** | **0.084** |
| Legume (%) | 0.135 | 0.096 | -0.133 | 0.088 | 0.033 | 0.088 |
| Woody (%) | -0.094 | 0.085 | 0.117 | 0.078 | -0.021 | 0.077 |
| Grass (%) | -0.107 | 0.092 | -0.012 | 0.085 | -0.098 | 0.083 |
| Non-native (%) | **-0.361** | **0.096** | -0.095 | 0.091 | **-0.254** | **0.088** |
| Soil trt. | -0.086 | 0.063 | -0.005 | 0.059 | 0.00 | 0.060 |
| Herb trt. | -0.032 | 0.063 | -0.09 | 0.06 | 0.042 | 0.061 |
| Legume × legume (%) | -0.185 | 0.112 | **-0.238** | **0.111** | 0.035 | 0.113 |
| Woody × woody (%) | -0.112 | 0.089 | **-0.213** | **0.089** | 0.02 | 0.090 |
| Grass × grass (%) | **0.251** | **0.077** | 0.058 | 0.078 | **0.205** | **0.078** |
| Forb × forb (%) | 0.060 | 0.073 | **0.158** | **0.074** | -0.046 | 0.075 |
| Non-native sp. × non-native<br>(%) | 0.032 | 0.108 | 0.014 | 0.105 | -0.021 | 0.108 |
| Marg. R<sup>2</sup>/Cond. R<sup>2</sup> | 0.208/0.266 |  | 0.189/0.220 |  | 0.161/0.191 |  |

Overall, we found that plant characteristics influenced partner fidelity (H2). We observed that individual plants from a species that hosted many bacterial antagonists or fungal mutualists (i.e., high normalised degree) exhibited lower interaction fidelity to those partners than individual plants from more specialised focal species (Fig 3b). We observed no significant effect of either the soil or herbivore treatment on any type of individual plant interaction fidelity (Fig. 3c-d).

Individual plant interaction fidelity to fungal mutualists was 1.19 times higher for legumes than non-legumes, irrespective of the percentage of legumes in the mesocosm (Fig. 4a iii and Table 1). Individual plant interaction fidelity to fungal antagonists, on the other hand, was 1.59 times higher if the plant was a legume and the only legume in the mesocosm than if the plant was a legume surrounded by legumes. There was no relationship between whether an individual plant was a legume and their fidelity to bacterial antagonists, nor was this lack of fidelity to bacterial antagonists significantly related to the percentage of legume species in the mesocosm.

**Fig. 4.**
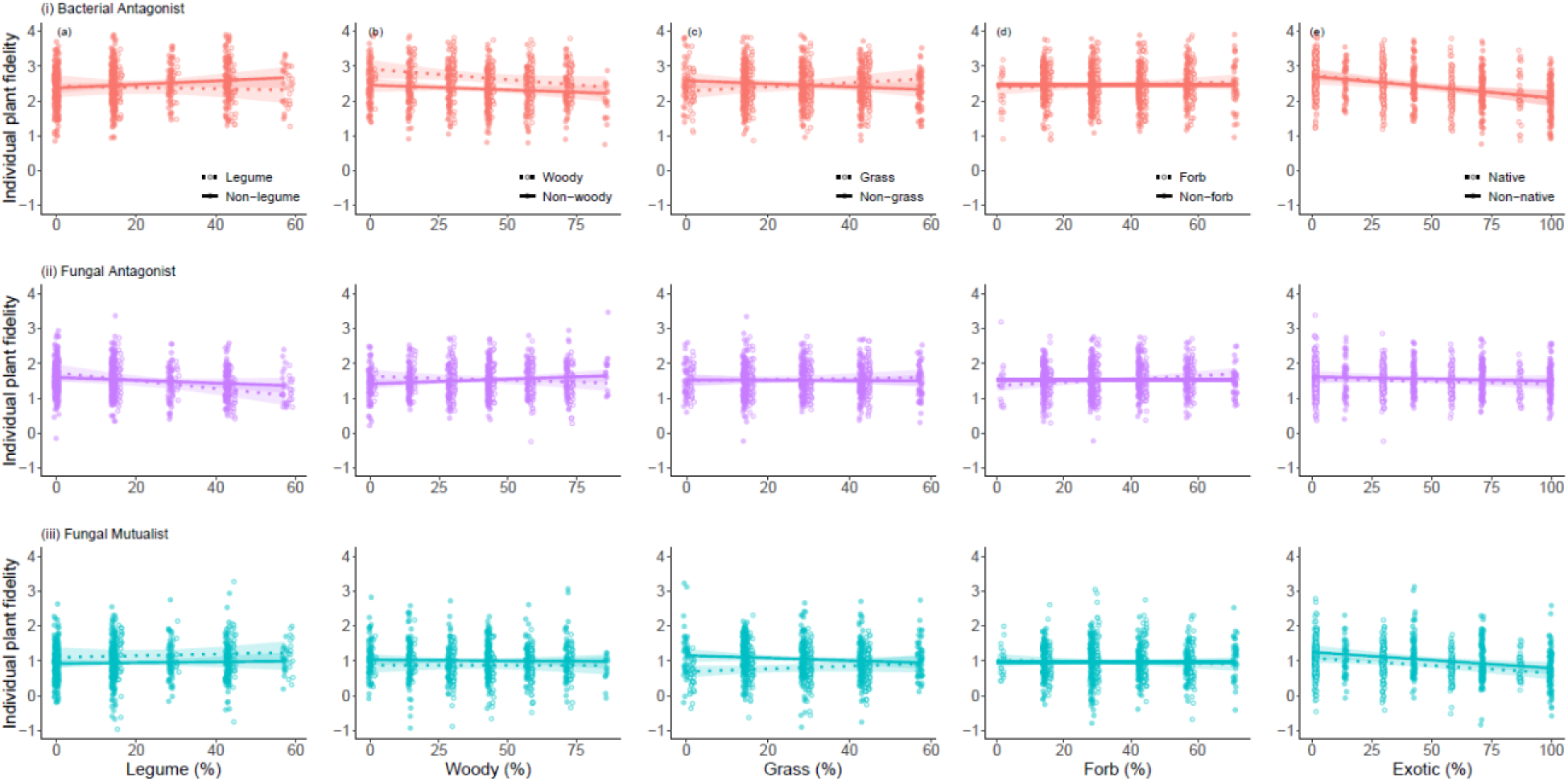
The inferred relationships between individual plant interaction fidelity to bacterial antagonists (red), fungal antagonists (purple), and fungal mutualists (turquoise) and the percentage of (a) legume species in the community for non-legumes (solid line) and legumes (dotted line); (b) woody species in the community for non-woody (solid line) and woody plants (dotted line); (c) grasses in the community for non-grasses (solid line) and grasses (dotted line); (d) forbs in the community for non-forbs (solid line) and forbs (dotted line); and (e) non-native species in the community for non-native plants (solid line) and native plants (dotted line) ± one standard deviation (shaded area). Further details including significance can be found in Table 1. Note that non-focal predictors were held at their mean values and all lines were fit using the linear mixed models described in the text.

Similar to the relationship between legume species surrounded by legume plants, woody plants had 1.13 times higher individual plant interaction fidelity to their fungal antagonists when there were no other woody plants in the mesocosm. On the other hand, there was no significant relationship between individual plant interaction fidelity to bacterial antagonists or fungal mutualists and the percentage of woody species in the community. However, woody species had 1.19 times higher individual plant interaction fidelity to their bacterial antagonists than non-woody species (Fig. 4bi; Table 1), whereas individual plant interaction fidelity to fungal mutualists was 1.18 times higher for non-woody species than woody species.

The impact of plant characteristics on partner fidelity (H2) also depended on the plant community context (H3). For example, there was a positive relationship between the percentage of grass species in a mesocosm and interaction fidelity for grasses to bacterial antagonists, and a negative relationship between the percentage of grass species in a mesocosm and interaction fidelity for non-grasses to bacterial antagonists (Fig. 4c ii; Table 1). There was a similar negative relationship between the percentage of grass species in a mesocosm and interaction fidelity for non-grasses to fungal mutualists, whereby non-grass fidelity to fungal mutualists was 1.22 times higher in mesocosms with the smallest percentage of grass plants. Interaction fidelity of grasses to fungal mutualists, on the other hand, had the opposite relationship (Fig. 4c iii). There was only a relationship between the percentage of forb species in a mesocosm and whether a plant was a forb for individual plant interaction fidelity to fungal antagonists whereby forbs in mesocosm communities with a high percentage of forbs had a higher fungal antagonist fidelity then a forb in a mesocosm with fewer forbs (Fig. 4d ii; Table 1).

Non-native plant species exhibited significantly higher individual interaction fidelity to their fungal mutualists than did native plants (1.16 times higher), irrespective of the percentage of non-native species in the community (Fig. 4e iii). Individual plant interaction fidelity to fungal mutualists did decline significantly, however, as the percentage of non-native species in the community increased. In fact, individual plant interaction fidelity to fungal mutualists was 1.58 times higher in mesocosms with the lowest percentage of non-native species.

The relationship between total plant biomass and individual plant interaction fidelity was context dependent (H4). However, this context dependency depended only on plant interaction fidelity to fungal mutualists and not to either type of belowground antagonist.

There was a significant interaction between the percentage of grass species in the mesocosm and individual plant interaction fidelity to fungal mutualists (Fig. 5a; Appendix S3 Table S2), irrespective of whether or not the focal plant was a grass. Specifically, a plant with low interaction fidelity to fungal mutualists was 54.2 times larger when there was a low percentage of grass species in the mesocosm, while a plant with high interaction fidelity to fungal mutualists was 22.3 times large when there was a high percentage of grass species in the mesocosm.

**Fig. 5.**
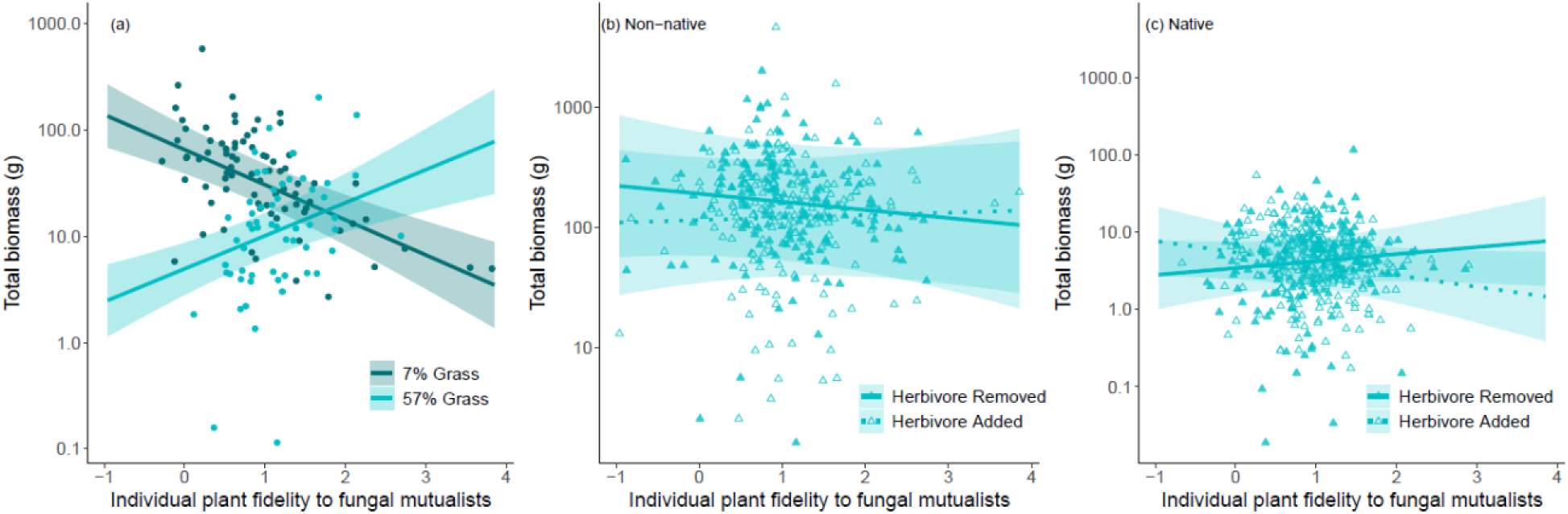
Inferred relationships between total biomass (grams on a logged scale) and individual plant interaction fidelity to fungal mutualists (turquoise) ± one standard deviation (shaded area). (a) The inferred relationship between total biomass and plant interaction fidelity to fungal mutualists where the gradation of lines represents the percentage of grass species in the mesocosm with the darker colour indicating fewer grass species, and the lighter colour more grass species. (b – c) The relationship between total biomass and plant interaction fidelity to fungal mutualists for (b) non-native plants and (c) native plants when herbivores are added (dashed lines) or removed (solid lines). All lines were fit using the linear mixed model described in the text.

Total plant biomass was significantly impacted by the three-way statistical interactions between the herbivory treatment, whether the plant was native or non-native, and individual plant interaction fidelity to its fungal mutualists. The total biomass of non-native plants decreased as individual plant interaction fidelity to fungal mutualists increased in mesocosms where herbivores were removed. The addition of herbivores, on the other hand, meant that high plant fidelity to fungal mutualists resulted in larger plants than low plant fidelity to fungal mutualists (Fig. 5e). This relationship was markedly different for native plants, whereby herbivores completely changed the relationship between individual plant interaction fidelity to fungal mutualists and total biomass. When herbivores were added to the mesocosms, native plant species were on average up to 5.2 times larger when they had low interaction fidelity to their fungal mutualists. When herbivores were removed, on the other hand, native plant species were on average up to 2.7 times larger when they had high interaction fidelity to their fungal mutualists.

The variables that explained total biomass were also related to belowground biomass (Appendix S3 Table S2). Unique to belowground biomass was a significant interaction between individual plant interaction fidelity to fungal antagonists and the percentage of non-native species in the mesocosm (Appendix S3 Fig. S1d -e; Table 2). Specifically, there was a positive relationship between interaction fidelity to fungal antagonists and non-native plant size when there were few other non-native plants in the mesocosm, with the reverse true when there a lot of non-native plants in the mesocosm. The converse was true for native plants.

## Discussion

Over the course of their lifetime, individual plants engage in a multitude of interactions, from beneficial interactions with mutualist partners to antagonistic interactions with harmful soil biota and herbivores. How plant fidelity to these interaction partners responds to a changing community context, and whether these responses relate to plant performance, have remained open questions. Our results demonstrate that individual plant interaction fidelity and its relationship to performance both vary depending on the type of interaction partner and community context being examined. In general, interactions between plants and their antagonist partners (both fungi and bacteria) were more conserved than expected by chance (H1), and, more highly conserved than interactions between plants and their mutualist partners. Furthermore, individual plant interaction fidelity to antagonist and mutualist partners depended on the characteristics of the plant (H2), such as functional group and provenance, and the community context (H3), such as the percentage of grass species in a community. Finally, individual plant performance was influenced primarily by interaction fidelity to fungal mutualist partners rather than antagonist partners (H4), with the strength of this relationship dependent on plant characteristics and community context.

Interestingly, we observed that individual plant fidelity to microbial interaction partners responded more strongly to indirect changes to the surrounding plant community and the consequences of plant-plant competition than to direct manipulations of the soil microbial community (the experimental home vs. away soil treatment). Moreover, this impact of the surrounding plant community on individual plant-microbe interactions depended strongly on focal plant type and the type of plants in the surrounding community (i.e., grass species in a mesocosm with a high percentage of grass species; Fig. 4, Table 1). Consequently, this suggests that functionally similar neighbours support the availability of specialist mutualists and antagonists. In this way, grasses had higher interaction partner fidelity when there was a high percentage of other grass species in the mesocosm supporting grass specialist mutualist and antagonists (Fig. 4). Interestingly, we only observed a significant interaction between community context and interaction fidelity on plant biomass—rather than a significant three-way interaction between type of plant, surrounding community, and interaction fidelity – demonstrating that the importance of partner fidelity is highly context dependent. That is, when the community has very few grass species, lower partner fidelity results in larger individuals than high partner fidelity. Notably, this was only significant for fungal mutualists and highlights the importance of considering interaction specificity rather than simply the number, or diversity, of interactions.

In general, we observed that individual plant interaction fidelity to antagonists was not related to plant biomass, despite the fact that individual plant interaction fidelity to antagonists was more predictable than individual plant interaction fidelity to fungal mutualists (Table 1). Interaction fidelity to antagonistic interaction partners can arise through multiple mechanisms. For example, plants can alter their microbiome to select for a ‘core microbiota’ (i.e., high interaction fidelity) through resource allocation or defensive compounds (McLaren & Callahan 2020; Trivedi *et al*. 2020). The cost of such defenses can limit resource allocation to growth (Zhou *et al*. 2022), such that a growth-defense trade-off could generate a negative correlation between partner fidelity and biomass. Pathogens can also interact with each other (Abdullah *et al*. 2017), such that a pathogen that is consistently successful on a given plant species and also able to consistently competitively exclude others (e.g., Al-Naimi *et al*. 2005), could generate a pattern of high plant interaction fidelity and impact plant biomass. While we observed a strong relationship between both plant characteristics and community characteristics on plant fidelity to antagonist partners, overall plant biomass was not affected by plant fidelity to antagonist partners (Table S2). This does not mean that antagonist partner fidelity is not important, however, it suggests that the importance is likely more nuanced with high partner fidelity to antagonists benefitting specific plants in specific circumstances, or dependent on total antagonist fidelity rather than specific fidelity to bacterial antagonists or fungal antagonists. Because research to date has typically focused on a single type of interaction partner (Porter *et al*. 2020), our understanding of the impact of these interdependencies on plant performance is still in its infancy, but our study takes a strong step forward by demonstrating the strong positive relationship between different types of fidelity and their differential impacts (or lack thereof) on plant performance.

Above- and belowground biomass were both associated the three-way interaction between whether the plant was native, whether herbivores were added, and individual plant interaction fidelity to fungal mutualists (Fig 5; Appendix S3 Table S2 and Fig S1). Notably, the significant interaction between aboveground herbivory and belowground mutualist interactions demonstrates that aboveground interactions can also impact belowground plant biomass (Bardgett & Wardle 2003; Wardle 2006). Furthermore, the nature of the relationship between plant biomass and herbivory depended on plant provenance – when herbivores were present, native species with low fidelity to their fungal mutualist partners were larger than those with high fidelity. It has been shown that generalist plants (i.e., plants with low fidelity) obtain complementary benefits through their diverse set of AMF interactions (Jansa *et al*. 2008; Koide 2000), including pathogen and stress resistance (Begum *et al*. 2019; Lutz *et al*. 2023), but similar to the results from our study, this generality comes with costs (Lewe *et al*. 2025). Interestingly, non-native species with high fidelity to fungal mutualists performed worse than those with low fidelity, irrespective of the presence of herbivores, perhaps reflecting their lack of coevolutionary history with native mutualists, or their reduced dependence on fungal mutualists compared with natives (Waller *et al*. 2016). One of the consequences of anthropogenic change is the formation of ecologically novel communities composed of species that may not have previously coexisted (Hobbs *et al*. 2006). Our study is an important step in addressing this question by experimentally manipulating functional traits and community contexts and exploring the impact individual responses to changes in biotic.

The composition of ecological communities is constantly changing, and individuals can either respond to or resist these changes. Previous studies on the net effect of species interactions on plant communities have highlighted strong context dependency (Chamberlain *et al*. 2014); however, here we demonstrate that plant interaction fidelity to belowground partners and its consequences for plant biomass depend on plant specific characteristics, and perhaps more intriguingly, the aboveground community context over multiple trophic levels (plants and herbivores). By manipulating broad biotic gradients, we were able to identify functional groups of plants that were better able to respond to being pushed beyond their ‘biotic comfort zone’ and to pinpoint the trade-offs in interaction fidelity with changing community context. In this way, our study represents a critical step in predicting plant performance in a changing world.

## Supporting information

All Supplemental Information

## Acknowledgements

This research was supported by funding from Bioprotection Aotearoa, a Centre of Research Excellence funded by the Tertiary Education Commission, New Zealand. AMM was supported by an NSERC postdoctoral fellowship and a Rutherford Discovery Fellowship administered by the Royal Society Te Apārangi (RDF-18-UOC-007 to JDT). JDT is supported by a Rutherford Discovery Fellowship administered by the Royal Society Te Apārangi (RDF-18-UOC-007).

JMT is funded by the Ministry of Business, Innovation & Employment (programme number C09X2209) and the Marsden Fund. BBM was support by a by a Rutherford Discovery Fellowship from New Zealand Government funding, which is managed by the Royal Society Te Aparangi (RDF-13-UOC-003 to DBS). DBS and HRL acknowledge the support from the Marsden Fund Council from New Zealand Government funding, which is managed by the Royal Society Te Aparangi (Grant 16-UOC-008 awarded to DBS).

