## Supplementary material for "Partner fidelity with root symbionts impacts plant performance in the face of changing above- and belowground community context": All Supplemental Information

**Supplemental material:**

**Appendix S1: In-depth Hypotheses**

**Appendix S2: Metric Description**

**Appendix S3: Additional Results**

**Appendix S1: In-depth Hypotheses**

In this appendix we provide additional details and examples underpinning the hypotheses described in the text.

(H1) We hypothesised that individual plant fidelity would depend on the type of interaction partner and be highest for fungal mutualists, given that interactions with mutualists are less evolutionarily malleable (Fortuna *et al.* 2020). Additionally, we hypothesised that individual plant fidelity to different types of interaction partners would be interdependent. It has been shown that the strength of different types of biotic interactions depend upon one another (Morris *et al.* 2007; Sauve *et al.* 2016). For example, flexibility of plants in their interactions with arbuscular mycorrhizal fungi (AMF) partners can result in a decrease in bacterial biomass in the rhizosphere (Lewe *et al.* 2025).

(H2) We hypothesised that plant characteristics would influence individual plant fidelity and that the direction of the effect would depend on the type of plant fidelity and plant characteristic. For example, non-native plant species should exhibit lower individual plant fidelity to their partners than native plant species (Coux *et al.* 2021; Ramana *et al.* 2023a), and legumes are expected to exhibit lower individual plant fidelity to their pathogen partners (Jack *et al.* 2019a). Because exotic plants are highly generalist (Aizen *et al.* 2008) and may be lacking locally co-evolved partners (Ramana *et al.* 2023b), they often interact with the most abundant partners available (Coux *et al.* 2021), or the most generalist partners (Waller *et al.* 2024). Therefore, we expected non-native plant species to exhibit lower individual plant fidelity to their partners than native species. Similarly, woody plant species host more specific mycorrhizal fungi than non-woody plant species, especially grasses (Polme *et al.* 2016; Sepp *et al.* 2019). Thus, we expected woody species to exhibit higher individual plant fidelity to their fungal mutualist partners than grasses or forbs. Finally, rhizobial bacteria can protect their legume hosts from microbial pathogens (Jack *et al.* 2019b), therefore we expected individual leguminous plants to exhibit lower individual plant fidelity to their antagonists than non-legumes.

(H3) We hypothesised that biotic context would also influence individual plant fidelity and that the effect would depend on the type of plant fidelity and plant characteristic, driven mostly by competition. For example, plant coexistence is strongly impacted by competition (Tilman 1982), for abiotic resources but also for belowground interactions (Lekberg *et al.* 2018), and the ability of a plant to outcompete other plants within the mesocosm for preferred partners should depend on biotic context (Emer *et al.* 2018). Additionally, competition with other plants has been shown to impacts the fitness of pathogens on their host plants (Lively *et al.* 1995) and the efficacy of host plant defenses (Seabloom *et al.* 2018). Similarly, herbivory reduces AMF colonisation and alpha diversity and increases phylogenetic overdispersion (Frew et al. 2024), suggesting that partner fidelity of plants should increase under herbivory as AMF compete for scarcer resources. Thus, we expected that individual plant fidelity to mutualists would decline in mesocosms that are increasingly dominated by stronger competitors, such as a plant growing in a mesocosm dominated by non-native species (Lankau 2012). Collectively, these above- and belowground interactions may modify the seemingly high partner fidelity observed for grasses and forbs grown individually without competitors or herbivores (Lewe *et al.* 2025).

(H4) We hypothesised that high fidelity to mutualists will be costly for individual plants and lead to reduced performance. For example, it has been shown that species invasions alter mutualistic interactions between native plant species (Aizen *et al.* 2008), thus we expected low individual plant fidelity to mutualists to decrease the detrimental impacts of species invasions . As mesocosms become increasingly dominated by non-native species, we expected native plants with lower individual plant fidelity to mutualists to have higher biomass than those with high individual plant fidelity to mutualists. Additionally, we expected plants with lower individual plant fidelity to antagonists to be better able to adapt to the competitive challenges that neighbour plants may exert (e.g. along increasingly competitive gradients) and thus be more tolerant of pressure from antagonists (Waller *et al.* 2024), resulting in higher biomass.

**Appendix S2: Metric Description**

In this appendix we describe the details of the metrics used in statistical analysis and provide a reference table (Table S1) with each of the plant species, their four-letter species code, their functional group, whether or not they are legumes, and their provenance.

**Species’ normalised degree:** a measure of generalism, defined for a given species as the number of partners which with a species interacts, adjusted by the total richness of interaction partners within that mesocosm. For each plant species, we therefore obtained three values of normalised degree, one for each partner type.

**Individual plant fidelity:** a focal individual’s average Sorensen similarity index in interaction partners calculated from pairwise comparisons to all other individual plants of a given species and then compared to a null model where interaction partners are chosen from the pool of species shared between the mesocosms being compared. In this way, if individual plant fidelity is less than –0.67 it can be considered significantly less faithful than expected by chance (where –0.67 is the lower boundary of the interquartile range), while if individual plant fidelity is greater than 0.67 it can be considered significantly more faithful than expected by chance (where 0.67 is the upper boundary of the interquartile range). There are three types of fidelity – individual plant fidelity to bacterial antagonists, individual plant fidelity to fungal antagonists, and individual plant fidelity to fungal mutualists.

**Plant characteristics:** plants were characterised according to their functional group – woody, grass, or forb – as well as whether or not they were legumes or non-native. While functional groups were mutually exclusive, whether they were legumes or non-native were not. In this way an individual plant could be a legume, non-native, and from any of the three functional groups (woody, grass, or forb). See Table S1 for individual species characteristics.

**Biotic context** (i.e. percentage woody, grasses, forbs, legumes, and non-native plants in the mesocosm): the percentage of individuals in the mesocosm, excluding the focal individual, which are woody/grass/forb/legume/non-native. In this way, there are five different biotic contexts.

**Herbivore treatment**: all mesocosms were caged and invertebrate herbivores were deliberately established in half of the mesocosms and actively removed from the other half (see (Allen *et al.* 2021) for full methods).

**Soil treatment:** we imposed a plant-soil feedback treatment, whereby mesocosms contained either ‘home’ soil (i.e., soil inocula containing organisms typically associated with the community of plants in the mesocosm) or ‘away’ soil (i.e., soil inocula containing organisms not typically associated with the plant community; see (Waller *et al.* 2020, 2024) for more details).

**Mesocosm community:** Mesocosm community was a number between 1 and 20, since there were 20 unique combinations of starting plant species replicated across soil and herbivore treatments.

**Table S1:** The four-letter code for each of the 36 plant species along with their functional group, whether they are legumes or not, and their provenance.

| Code | Plant species | Functional group | Leguminous Sp. | Provenance |
| --- | --- | --- | --- | --- |
| AcCa | *Acaena caesiiglauca* | Forb | No | Native |
| AcDe | *Acacia dealbata* | Woody | Yes | Native |
| AcPu | *Acaena purpurea* | Forb | No | Native |
| AcMi | *Achillea millefolium* | Forb | No | Non-native |
| AgCa | *Agrostis capillaris* | Grass | No | Non-native |
| AnLe | *Anemanthele lessoniana* | Grass | No | Native |
| AnOd | *Anthoxanthum odoratum* | Grass | No | Non-native |
| BrGr | *Brachyglottis greyi* | Shrub | No | Native |
| CaSe | *Carex secta* | Grass | No | Native |
| CiVu | *Cirsium vulgare* | Forb | No | Non-native |
| CoRo | *Coprosma robusta* | Shrub | No | Native |
| DaGl | *Dactylis glomerata* | Grass | No | Non-native |
| EcVu | *Echium vulgare* | Forb | No | Non-native |
| FeNo | *Festuca novae-zelandiae* | Grass | No | Native |
| HoLa | *Holcus lanatus* | Grass | No | Non-native |
| HyPe | *Hypericum perforatum* | Forb | No | Non-native |
| LeSc | *Leptospermum scoparium* | Shrub | No | Native |
| LoPe | *Lolium perenne* | Grass | No | Non-native |
| LuAr | *Lupinus arboreus* | Forb | Yes | Non-native |
| MeSa | *Medicago sativa* | Forb | Yes | Non-native |
| MuAs | *Muehlenbeckia astonii* | Shrub | No | Native |
| MuCo | *Muehlenbeckia complexa* | Shrub | No | Native |
| OlVi | *Olearia virgata* | Shrub | No | Native |
| OzLe | *Ozothamnus leptophyllus* | Shrub | No | Native |
| PhCo | *Phormium cookianum* | Forb | No | Native |
| PiCo | *Pinus contorta* | Woody | No | Non-native |
| PiRa | *Pinus radiata* | Woody | No | Non-native |
| PoCi | *Poa cita* | Grass | No | Native |
| PoCo | *Poa colensoi* | Grass | No | Native |
| PoTo | *Podocarpus totara* | Woody | No | Native |
| RuAc | *Rumex acetosella* | Forb | No | Non-native |
| RuOb | *Rumex obtusifolius* | Forb | No | Non-native |
| SoMi | *Sophora microphylla* | Shrub | Yes | Native |
| TrPr | *Trifolium pratense* | Forb | Yes | Non-native |
| UlEu | *Ulex europaeus* | Shrub | Yes | Non-native |
| VeOd | *Veronica odora* | Shrub | No | Native |

**Appendix S3: Additional Results**

In this appendix we provide additional results including a full table of results from an analysis of how individual plant fidelity interacts with species characteristics and biotic community contexts to impact both total and belowground plant biomass (Table S2), and the inferred relationships between statistically significant predictors and belowground plant biomass (Fig. S1).

**Table S2.** The results from an analysis of how individual plant fidelity interacts with species characteristics and mesocosm contexts to impact total plant biomass and belowground plant biomass. The main effects of the model are presented first, followed by the interaction between individual plant fidelity to each partner type and each main effect. Here, BA represents bacterial antagonist, FA represents fungal antagonist, and FM represents fungal mutualist, Fid. is fidelity, sp. is species, Legume sp. is whether or not the focal plant is a legume, Woody sp. is whether or not the focal plant is woody, Grass sp. is whether or not the focal species is a grass, Non-native sp. represents whether the focal species is non-native, Soil trt. is the soil treatment (home or away), Herb trt. is the herbivore treatment (added or removed), Woody (%) is the percentage of woody species in that mesocosm (not counting the focal plant), and the same interpretation applies for Grass (%), Legume (%), and Non-native (%). Bold values represent significant relationships. Marg. R^2^ and Cond. R^2^ represent marginal and conditional R^2^, respectively where conditional R^2^ accounts for random effects. There is no forb species or forb (%) in the table because they correspond to situations when species are neither woody nor grasses.

|  |  | *Total* | | *Belowground* | |
| --- | --- | --- | --- | --- | --- |
|  |  | *Est.* | *Std. Error* | *Est.* | *Std. Error* |
| Main Effects | Fid. to BA | 0.061 | 0.169 | 0.076 | 0.179 |
|  | Fid. to FA | 0.016 | 0.148 | 0.005 | 0.156 |
|  | Fid. to FM | 0.097 | 0.154 | 0.123 | 0.164 |
|  | BA sp. degree | 0.109 | 0.376 | 0.488 | 0.301 |
|  | FA sp. degree | -0.491 | 0.419 | -0.337 | 0.334 |
|  | FM sp. degree | 0.138 | 0.356 | 0.161 | 0.284 |
|  | Legume sp. | -0.013 | 1.05 | 0.011 | 0.845 |
|  | Woody sp. | 0.858 | 0.599 | -0.598 | 0.483 |
|  | Grass sp. | **3.331** | **0.778** | 0.519 | 0.625 |
|  | Non-native sp. | **3.656** | **0.646** | **2.298** | **0.522** |
|  | Legume (%) | -0.155 | 0.22 | -0.165 | 0.21 |
|  | Woody (%) | 0.298 | 0.18 | **0.388** | **0.175** |
|  | Grass (%) | -0.279 | 0.193 | -0.217 | 0.186 |
|  | Non-native (%) | **-0.466** | **0.207** | -0.38 | 0.202 |
|  | Soil trt. | -0.014 | 0.078 | 0.049 | 0.083 |
|  | Herb trt. | -0.083 | 0.104 | -0.076 | 0.111 |
|  | Legume sp. × Legume (%) | -0.322 | 0.22 | -0.186 | 0.232 |
|  | Woody sp. × Woody (%) | 0.027 | 0.156 | -0.113 | 0.166 |
|  | Grass sp. × Grass (%) | -0.069 | 0.127 | -0.028 | 0.134 |
|  | Forb sp. × Forb (%) | -0.082 | 0.118 | -0.006 | 0.126 |
|  | Non-native sp. × Non-native (%) | -0.003 | 0.196 | -0.225 | 0.209 |
|  | Provenance × Herb | -0.221 | 0.168 | -0.235 | 0.178 |
| Interactions between each predictor and fidelity | |  |  |  |  |
| Fid. to FA | Fid. to BA | -0.051 | 0.05 | -0.049 | 0.053 |
| Fid. to FA | Fid. to FM | -0.021 | 0.049 | 0.034 | 0.052 |
| Fid. to FM | Fid. to BA | 0.058 | 0.053 | 0.037 | 0.057 |
| BA Sp. degree | Fid. to BA | 0.013 | 0.082 | -0.053 | 0.087 |
| FA Sp. degree | Fid. to FA | 0.028 | 0.064 | -0.013 | 0.067 |
| FM Sp. degree | Fid. to FM | -0.046 | 0.054 | -0.008 | 0.057 |
| Legume sp. | Fid.to BA | 0.254 | 0.241 | 0.353 | 0.255 |
|  | Fid. to FA | 0.204 | 0.201 | 0.224 | 0.213 |
|  | Fid. to FM | -0.141 | 0.193 | -0.096 | 0.204 |
| Woody sp. | Fid.to BA | -0.045 | 0.157 | -0.134 | 0.166 |
|  | Fid. to FA | -0.153 | 0.138 | -0.161 | 0.146 |
|  | Fid. to FM | 0.227 | 0.145 | 0.241 | 0.153 |
| Grass sp. | Fid.to BA | 0.028 | 0.174 | 0.004 | 0.184 |
|  | Fid. to FA | 0.001 | 0.149 | 0.054 | 0.157 |
|  | Fid. to FM | -0.035 | 0.142 | -0.025 | 0.15 |
| Non-native sp. | Fid.to BA | -0.207 | 0.188 | -0.143 | 0.199 |
|  | Fid. to FA | 0.103 | 0.159 | 0.187 | 0.169 |
|  | Fid. to FM | -0.21 | 0.174 | **-0.386** | **0.185** |
| Legume (%) | Fid.to BA | 0.033 | 0.082 | 0.108 | 0.087 |
|  | Fid. to FA | 0.011 | 0.079 | -0.054 | 0.084 |
|  | Fid. to FM | 0.092 | 0.085 | 0.127 | 0.091 |
| Woody (%) | Fid.to BA | -0.054 | 0.086 | -0.08 | 0.091 |
|  | Fid. to FA | -0.052 | 0.087 | -0.061 | 0.092 |
|  | Fid. to FM | 0.149 | 0.1 | 0.138 | 0.106 |
| Grass (%) | Fid.to BA | 0.016 | 0.089 | 0.099 | 0.094 |
|  | Fid. to FA | -0.093 | 0.09 | -0.114 | 0.095 |
|  | Fid. to FM | **0.275** | **0.1** | **0.289** | **0.106** |
| Non-native (%) | Fid.to BA | -0.029 | 0.091 | -0.066 | 0.096 |
|  | Fid. to FA | 0 | 0.099 | 0.096 | 0.105 |
|  | Fid. to FM | -0.095 | 0.115 | -0.041 | 0.122 |
| Soil trt. | Fid.to BA | 0.043 | 0.085 | 0.034 | 0.09 |
|  | Fid. to FA | 0.127 | 0.087 | 0.049 | 0.092 |
|  | Fid. to FM | -0.044 | 0.085 | 0.032 | 0.09 |
| Herb trt. | Fid.to BA | -0.044 | 0.11 | -0.029 | 0.117 |
|  | Fid. to FA | 0.005 | 0.116 | 0.084 | 0.123 |
|  | Fid. to FM | **-0.32** | **0.124** | **-0.289** | **0.132** |
| Non-native sp. | Fid.to BA | 0.047 | 0.18 | 0.217 | 0.191 |
| × Herb | Fid. to FA | -0.26 | 0.178 | -0.354 | 0.189 |
|  | Fid. to FM | **0.438** | **0.173** | **0.418** | **0.183** |
| Legume sp. × | Fid.to BA | -0.072 | 0.185 | -0.1 | 0.195 |
| Legume (%) | Fid. to FA | -0.25 | 0.179 | -0.207 | 0.189 |
|  | Fid. to FM | 0.09 | 0.137 | 0.214 | 0.145 |
| Woody sp. × | Fid.to BA | -0.062 | 0.119 | -0.008 | 0.126 |
| Woody (%) | Fid. to FA | 0.047 | 0.12 | 0.043 | 0.127 |
|  | Fid. to FM | -0.104 | 0.131 | -0.084 | 0.138 |
| Grass sp. × | Fid.to BA | 0.16 | 0.109 | 0.147 | 0.115 |
| Grass (%) | Fid. to FA | -0.11 | 0.108 | -0.026 | 0.114 |
|  | Fid. to FM | -0.113 | 0.126 | -0.13 | 0.133 |
| Forb sp. × | Fid.to BA | -0.117 | 0.113 | -0.148 | 0.12 |
| Forb (%) | Fid. to FA | -0.144 | 0.105 | -0.093 | 0.111 |
|  | Fid. to FM | 0.127 | 0.108 | 0.206 | 0.114 |
| Non-native sp. | Fid.to BA | 0.044 | 0.126 | 0.001 | 0.134 |
| × Non-native | Fid. to FA | -0.192 | 0.121 | **-0.268** | **0.128** |
| (%) | Fid. to FM | 0.225 | 0.136 | 0.132 | 0.144 |
| Marg. R^2^/Cond. R^2^ | | 0.381/0.863 | | 0.280/0.759 | |

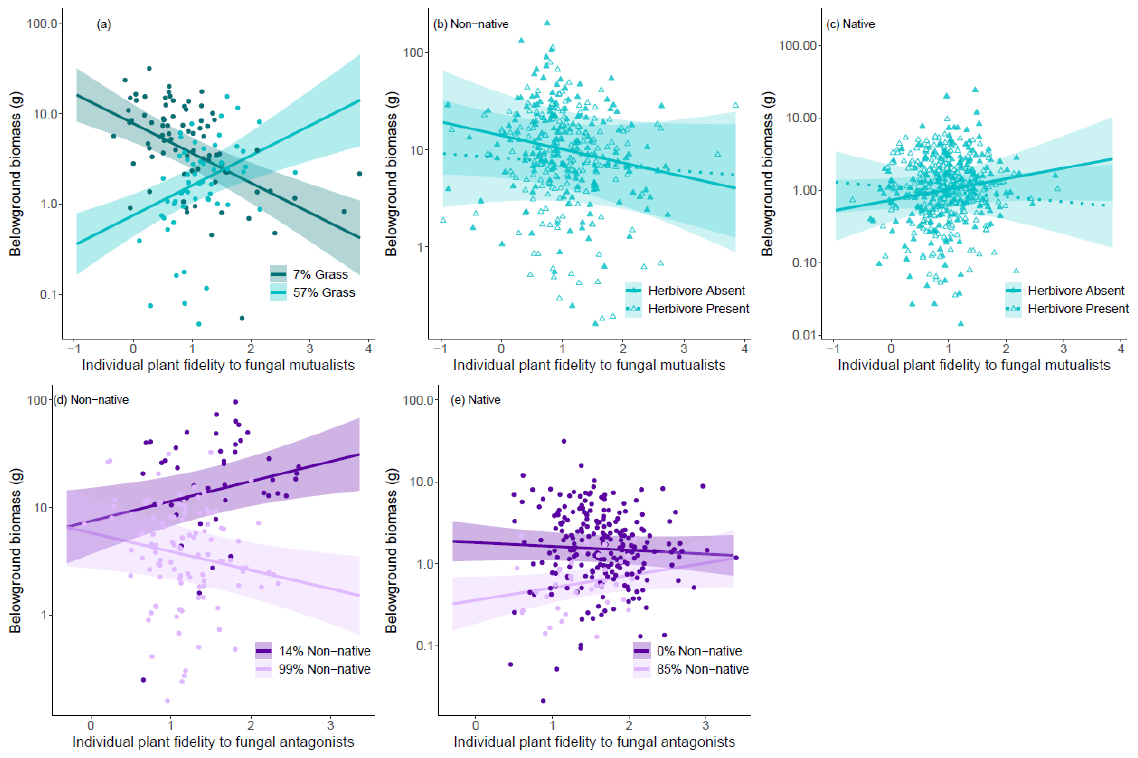

**Fig. S1**. Inferred relationships between belowground biomass (g) and individual plant fidelity to fungal antagonists (purple), or fungal mutualists (turquoise), ± one standard deviation (shaded area). (a) The inferred relationship between belowground biomass and plant interaction fidelity to fungal mutualists where the gradation of lines represents the percentage of grass species in the mesocosm with the darker colour indicating fewer grass species, and the lighter colour more grass species. (b – c) The relationship between belowground biomass and plant interaction fidelity to fungal mutualists for (b) non-native plants and (c) native plants when herbivores are added (dashed lines) or removed (solid lines). (d – e) The relationship between belowground biomass and plant interaction fidelity to fungal antagonists for (d) non-native plants and (e) native plants where the gradation of lines represents the percentage of non-native plants in the mesocosm with the darker colour indicating fewer non-native species and the lighter colour more non-native species. All lines were fit using the linear mixed model described in the text.
